# Insula Functional Connectivity in Schizophrenia

**DOI:** 10.1101/2019.12.16.878827

**Authors:** Julia M. Sheffield, Baxter P. Rogers, Jennifer Urbano Blackford, Stephan Heckers, Neil D. Woodward

## Abstract

The insula is structurally abnormal in schizophrenia, demonstrating robust reductions in gray matter volume, cortical thickness, and altered gyrification during prodromal, early and chronic stages of the illness. Despite compelling structural alterations, less is known about its functional connectivity, limited by studies considering the insula as a whole or only within the context of resting-state networks. There is evidence, however, from healthy subjects that the insula is comprised of sub-regions with distinct functional profiles, with dorsal anterior insula (dAI) involved in cognitive processing, ventral anterior insula (vAI) involved in affective processing, and posterior insula (PI) involved in somatosensory processing. The current study builds on this prior work and characterizes insula resting-state functional connectivity sub-region profiles in a large cohort of schizophrenia (N=191) and healthy (N=196) participants and hypothesizes specific associations between insula sub-region connectivity abnormalities and clinical characteristics related to their functional profiles. Functional dysconnectivity of the insula in schizophrenia is broadly characterized by reduced connectivity within insula sub-networks and hyper-connectivity with regions not normally connected with that sub-region, reflected in significantly greater similarity of dAI and PI connectivity profiles and significantly lower similarity of dAI and vAI connectivity profiles (p<.05). In schizophrenia, hypo-connectivity of dAI correlates with cognitive function (r=.18, p=.014), whereas hyper-connectivity between vAI and superior temporal sulcus correlates with negative symptoms (r=.27, p<.001). These findings reveal altered insula connectivity in all three sub-regions and converges with recent evidence of reduced differentiation of insula connectivity in schizophrenia, implicating functional dysconnectivity of the insula in cognitive and clinical symptoms.

## 1. Introduction

Abnormal structure of the insula is one of the most robust anatomical findings in schizophrenia. Meta-analyses have concluded that grey matter volume is reduced in bilateral insula and that smaller insula volume, while present in other mental illnesses, is most pronounced in psychotic disorders (Goodkind et al., 2015; Shepherd et al., 2012). Insula volume alterations are seen across the disease course (Ellison-Wright et al., 2008), demonstrating reductions in first-episode patients within the first few years of illness (Lee et al., 2016) with even more robust, progressive reductions during the chronic state (Chan et al., 2011). Neurobiological alterations of the insula may even represent a useful early marker of schizophrenia-risk, as smaller initial bilateral insula volume is seen in ultra-high risk patients who transition to psychosis, as compared to ultra-high risk patients who do not transition (Takahashi et al., 2009). Combined, these findings implicate insula cortex integrity in the pathophysiology of schizophrenia.

Considerably less is known about functional abnormalities of the insula in schizophrenia. This critical knowledge gap is likely due to the fact that the insula is involved in a diverse range of functions (Wylie and Tregellas, 2010), including cognition (e.g. cognitive control, error processing), social and emotion processing (e.g. disgust), interoception, and pain (Craig, 2009; Namkung et al., 2017; Uddin, 2015). The functional diversity of the insula is reflected in its heterogeneous structure and connectivity. At the level of cytoarchitecture, there are at least three sub-divisions of the insula, including a dysgranular dorsal anterior region (dAI), an agranular ventral anterior region (vAI), and a granular posterior region (PI) (Mesulam and Mufson, 1982). Identification and specificity of these sub-regions in humans has been confirmed through convergent analysis of task-based activations and connectivity assessed with functional neuroimaging (Deen et al., 2011; Nelson et al., 2010). Meta-analysis of functional co-activation during performance of various tasks has revealed preferential involvement of the dAI in cognition, the vAI in affective processing, and the PI in somatosensory processes (Uddin et al., 2013). The different functional roles of insula sub-regions are recapitulated by their dissociable functional connectivity profiles. The dAI is functionally connected to regions associated with higher-level cognition, including the dorsal anterior cingulate cortex (dACC), prefrontal cortex, frontal eye fields and intraparietal sulcus (Deen et al., 2011). Consistent with its role in emotion, the vAI exhibits robust functional connectivity with other emotion-related brain areas, including the pregenual anterior cingulate, inferior frontal gyrus, portions of the superior temporal sulcus (Deen et al., 2011), and the amygdala (Mutschler et al., 2009). Similarly, the PI, which is involved in sensorimotor processing, is connected to primary and secondary somatosensory and motor cortices (Deen et al., 2011). The insula is also highly interconnected with itself, underscoring its role in integrating cognitive, emotion, and sensorimotor information (Deen et al., 2011; Uddin et al., 2013).

Studying the functional connectivity of insula sub-regions in schizophrenia may expand our understanding of the pathophysiology of the disorder and inform the neural substrate of clinical phenotypes (Wylie and Tregellas, 2010). However, the majority of studies looking at the insula in schizophrenia investigated either connectivity of the whole insula (Pang et al., 2017) or within the context of broader cortical resting-state networks that include the insula, such as the salience, cingulo-opercular, and somatosensory networks (Huang et al., 2019; Sheffield et al., 2015). Directed connectivity of the anterior insula (a hub of the salience network) to nodes within the default mode network (DMN) and central executive network is reduced in schizophrenia (Moran et al., 2013; Palaniyappan et al., 2013). Salience network connectivity can also predict diagnostic group in first-episode patients and healthy controls with significant accuracy (Mikolas et al., 2016). Studies investigating connectivity of the insula with the whole brain have observed a range of findings in relatively small samples of schizophrenia participants (N<50) including: reduced connectivity of the whole insula with the ACC, caudate, and Heschl’s gyrus (Pang et al., 2017), increased connectivity between right PI and thalamus (Chen et al., 2016), and reduced connectivity of the vAI with regions of the DMN (Moran et al., 2013). Recent analysis also revealed reduced differentiation of insula connectivity sub-networks for anterior and posterior regions (Tian et al., 2019). Together, these studies indicate insula functional connectivity alterations in schizophrenia that have yet to be fully characterized in a large sample of patients.

The current study leverages a large resting-state fMRI dataset to characterize functional connectivity of insula sub-regions in schizophrenia and test specific *a priori* hypotheses about the associations between dysconnectivity of specific insula sub-regions and clinical phenotypes. Given the dAI’s well-defined role in higher-order cognition (Dosenbach et al., 2006; Moran et al., 2013), we hypothesized that dAI connectivity would be reduced in schizophrenia and associated with impaired general cognitive ability. The vAI is implicated in emotion processing (Simmons et al., 2013), a core feature of social cognition believed to underlie negative symptoms (Kohler et al., 2000; Martin et al., 2005). Therefore, we hypothesized that reduced vAI connectivity with affective regions would contribute to negative symptom severity. Finally, the PI plays a critical role in interoception, perception, and sensorimotor integration (Craig, 2009; Cauda et al., 2012), aspects of the human experience that are altered in psychosis. Greater interoceptive awareness in schizophrenia has been associated with more severe psychosis (Ardizzi et al., 2016) and a region in the insula that was activated during a symptom-capture study of auditory hallucinations was hyper-connected to the cerebellum and angular gyrus when compared to controls (Mallikarjun et al., 2018). We therefore hypothesized that hyper-connectivity of the PI would be associated with more severe positive symptoms.

## 2. Materials and methods

### 2.1 Participants

210 healthy individuals and 234 people with a schizophrenia spectrum disorder (i.e. schizophrenia, schizophreniform disorder, or schizoaffective disorder-hereafter referred to as “schizophrenia”) participated in one of three MRI studies (CT00762866; 1R01MH070560; 1R01MH102266) conducted in the Department of Psychiatry and Behavioral Sciences at Vanderbilt University Medical Center (VUMC) (include/exclusion criteria described in Supplement). Schizophrenia participants were recruited from the Psychotic Disorders Program at VUMC. Healthy controls were recruited from Nashville and the surrounding area via advertisements and word of mouth. The study was approved by the Vanderbilt Institutional Review Board and written informed consent was obtained before participation. The Structured Clinical Interview for DSM-IV (SCID-IV) was administered to all study participants to confirm diagnoses in people with schizophrenia and rule out psychopathology in healthy individuals (First et al., 1995). From our initial cohort of 444 individuals, 14 healthy and 43 schizophrenia participants were excluded for data quality reasons (described in Supplement). Thus, the final sample included 196 healthy and 191 schizophrenia participants (Table 1).

**Table 1.**
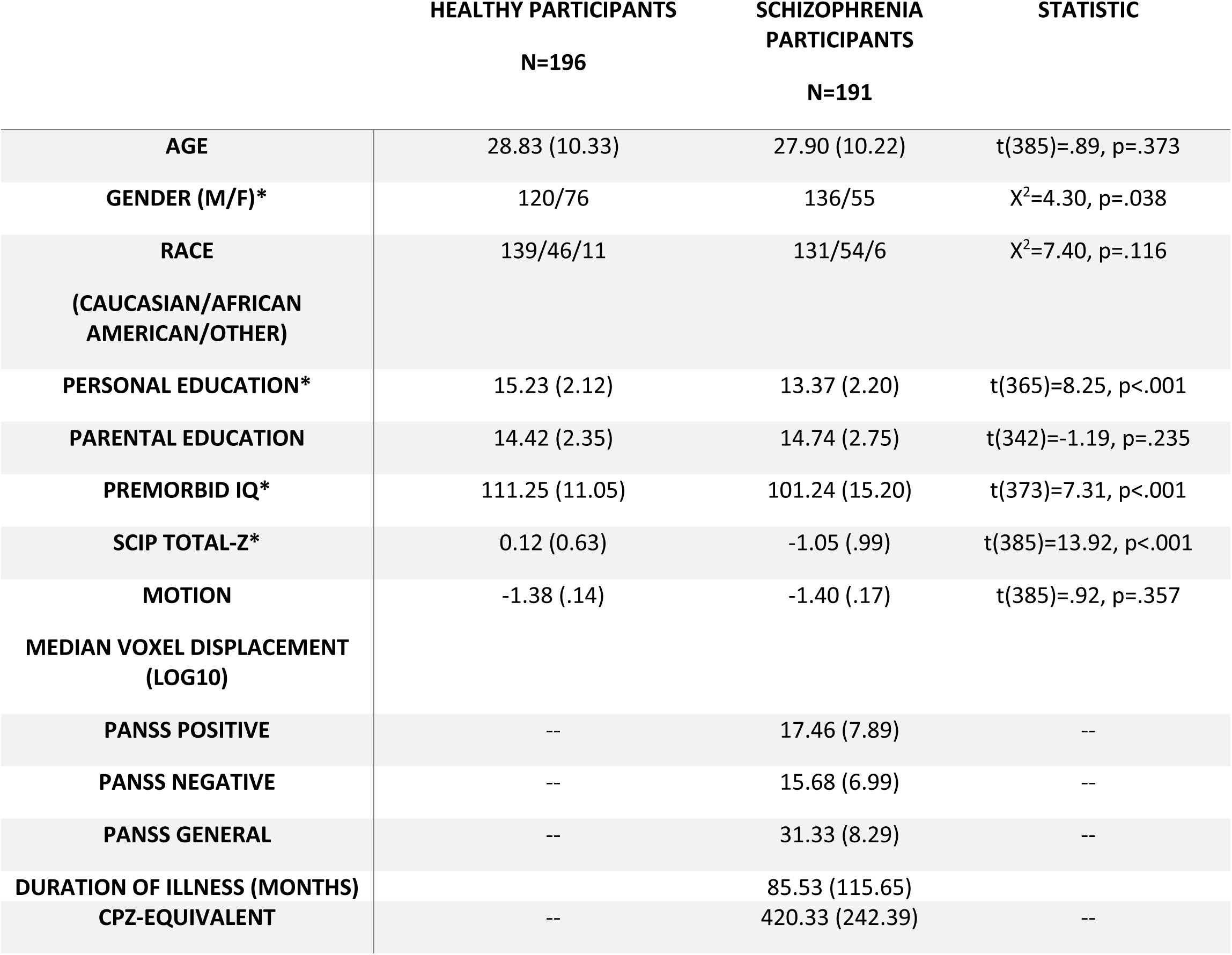

### 2.2 Clinical Data and Cognitive Data

Positive and negative symptoms of psychosis were quantified with the Positive and Negative Syndrome Scale (PANSS (Kay et al., 1987)). One subject was missing a positive scale score, so the average PANSS positive score across all schizophrenia participants was used for that subject. All individuals were administered the Screen for Cognitive Impairment in Psychiatry (SCIP), a brief (10-15 minute) measure of immediate and delayed verbal memory, verbal fluency, working memory and processing speed (for description of SCIP see (Menkes et al., 2018)). Scores from each sub-test were z-scored based on published norms (Purdon, 2005) and summed as a measure of cognitive ability.

### 2.3 Neuroimaging Data Acquisition and Preprocessing

Neuroimaging data were collected on one of two identical 3T Philips Intera Achieva scanners located at Vanderbilt University Institute of Imaging Sciences. Details of scanning parameters and pre-processing steps are included in the Supplement. Briefly, a seven or ten-minute echo-planar imaging (EPI) resting-state fMRI scan and T1-weighted anatomical (1mm isotropic resolution) were collected for each participant. Preprocessing of fMRI data, performed in SPM12, included correction for head motion using rigid body motion correction, spatial co-registration to T1-weighted anatomical images, and spatial normalization to MNI-space using the parameters obtained from the grey matter segmentation normalization.

### 2.4 Functional Connectivity of Insula Sub-Regions

A-priori masks for left and right insula sub-regions (dorsal anterior insula (dAI), ventral anterior insula (vAI), and posterior insula (PI)), derived by Deen and colleagues ((Deen et al., 2011) https://bendeen.com/data/) were used as seeds in separate seed-to-voxel functional connectivity analyses in MNI-space. Mean gray matter signal, six head motion parameters and their first temporal derivatives, and six principal components from an eroded white matter/CSF mask obtained from the individual T1 segmentations were removed in voxelwise regression (Behzadi et al., 2007). The fMRI time-series data was simultaneously band-pass filtered (0.01-0.10 Hz). Time series were averaged from voxels within each insula ROI and correlated with every voxel to generate a whole-brain connectivity map for each insula sub-region. Correlations were converted to Fisher Z-scores.

As the goal of the current project was to characterize insula connectivity in schizophrenia without specific hypotheses regarding laterality, the resultant functional connectivity maps for insula sub-regions were averaged across hemisphere to generate average connectivity maps for right/left sub-regions (results for left/right insula sub-regions presented in Table S1). The insula sub-region connectivity maps were smoothed using 4mm FWHM Gaussian kernel.

### 2.5 Statistical Analyses

Voxel-wise analyses of neuroimaging data were done in SPM12. Prior to comparing insula connectivity between groups, the connectivity profiles of each insula sub-region were qualitatively examined separately in the healthy and schizophrenia groups using one-sample t-tests thresholded at whole-brain (FWE) cluster-level corrected p=.05 using a voxel-wise p=.001 (uncorrected) cluster-defining threshold.

To compare connectivity between groups, insula sub-region connectivity maps were entered into separate independent-sample t-tests, controlling for gender and the research study for which the data was collected (given the three studies noted above). Independent samples t-tests were thresholded at cluster-level p_FWE_<.05 for voxel-wise cluster-defining threshold p=.001 (uncorrected). Connectivity results from the healthy>schizophrenia contrast are discussed as “hypo-connectivity” and results from the schizophrenia>healthy contrast are discussed as “hyper-connectivity”. To help determine whether medication dose was driving group difference results, connectivity from clusters showing a significant group difference were correlated with chlorpromazine-equivalent (CPZ) values.

In addition to comparing connectivity between groups, we examined how similar/dissimilar insula sub-region connectivity profiles were to one another and compared this across groups using eta-squared (Cohen et al., 2008). This analysis provided additional insight into whether differentiation of sub-region functional profiles was reduced in schizophrenia. Eta-squared for dAI-PI, dAI-vAI, and PI-vAI within-group estimated the proportion of variance in one map (e.g. dAI) accounted for by variance in another map (e.g. PI).

Meta-analysis has revealed distinct connectivity profiles for insula sub-regions that suggest involvement of each sub-region in specific aspects of information processing (Uddin et al., 2013), motivating our a-priori brain-behavior hypotheses. To test these hypotheses, two strategies were used. First, to maximize statistical power and limit the number of correlations performed, functional connectivity was extracted from the clusters demonstrating group differences using the volume of interest (VOI) tool in SPM, resulting in average hypo- and/or hyper-connectivity measure of each insula seed for each individual. These values served as dependent variables in the correlation analyses, that were Bonferroni-corrected for three tests (.05/3 = significance threshold of p<.0167). We complemented this approach with whole-brain voxel-wise correlation analyses with the hypothesized clinical variable entered as a predictor of functional connectivity including study and head motion (median voxel-displacement log10) as covariates. Whole-brain correlation analyses were cluster-level corrected at p_FWE_ <.05 for voxel-wise cluster-defining threshold of p=.001 (uncorrected).

## 3. Results

### 3.1 Functional Connectivity of Insula Sub-regions

Insula sub-regions demonstrated relatively distinct patterns of connectivity that were qualitatively similar in healthy controls and schizophrenia, and highly consistent with prior studies of insula connectivity (e.g. (Deen et al., 2011); Fig.1). Briefly, dAI was connected with fronto-parietal regions, dACC, and left temporal sulcus. Connectivity with vAI was seen in ventral ACC, inferior parietal lobe, and inferior temporal sulcus. The PI was largely connected with primary sensory and motor cortices, as well as the anterior cingulate. All insula sub-regions were negatively connected with the regions in the DMN (i.e. precuneus, posterior cingulate cortex (PCC), medial prefrontal cortex, and posterior cerebellum).

### 3.2 Insula Connectivity Abnormalities in Schizophrenia

Group differences in insula connectivity are shown in Fig. 2 and summarized in Table 2. Group differences were highly consistent across both right and left insula and were largely unrelated to CPZ-equivalent doses (Table S2)

**Table 2:** Connectivity differences between healthy and schizophrenia participants for insula sub-regions

| Seed Region of Interest, Contrast, and Brain Regions | Montreal Neurological Coordinates (x,y,z) | Brodmann Area | Peak t-value | p<sup>a</sup> | Cluster Size (Voxels) |
| --- | --- | --- | --- | --- | --- |
| **Dorsal Anterior Insula** |  |  |  |  |  |
| *Healthy Controls > Schizophrenia* |  |  |  |  |  |
| Left Substantia Innominata | -20, 8, -10 | 49 | 5.76 | <.001 | 487 |
| Right Substantia Innominata | 22, 2, -12 | 49 | 5.13 | <.001 | 441 |
| Right Dorsal Anterior Cingulate Cortex (dACC) | 2, 38, 6 | 24/32 | 4.70 | 0.029 | 135 |
| Left Orbitofrontal Cortex (OFC) | -30, 36, -8 | 47 | 4.26 | 0.004 | 207 |
| *Schizophrenia > Healthy Controls* |  |  |  |  |  |
| Left Superior Parietal Lobe | -24 -54 60 | 7 | 5.35 | <.001 | 1117 |
| Right Somatosensory Cortex | 42, -32, 58 | 1 | 4.36 | <.001 | 985 |
| Cerebellum (Ventral Attention Network)<sup>b</sup> | 40, -52, -32 | N/A | 4.79 | 0.039 | 126 |
| Cerebellum (Ventral Attention Network)<sup>b</sup> | 26, -70, -20 | N/A | 4.23 | <.001 | 290 |
| Primary Visual Cortex | 22, -68, 10 | 17 | 3.98 | 0.001 | 261 |
| **Ventral Anterior Insula** |  |  |  |  |  |
| *Healthy Controls > Schizophrenia* |  |  |  |  |  |
| Right Dorsal Anterior Cingulate Cortex (dACC) | 2, 38, 6 | 24/32 | 5.74 | <.001 | 591 |
| *Schizophrenia > Healthy Controls* |  |  |  |  |  |
| Cerebellum (Default Mode Network)<sup>b</sup> | 40, -60, -38 | N/A | 4.37 | 0.038 | 124 |
| Precuneus | -8, -54, 52 | 7 | 3.81 | 0.041 | 122 |
| **Posterior Insula** |  |  |  |  |  |
| *Healthy Controls > Schizophrenia* |  |  |  |  |  |
| Left Hippocampus | -24, -10, -12 | 54 | 4.54 | 0.025 | 129 |
| Superior Temporal Gyrus (STG) | -56, -6, -10 | 22 | 4.22 | 0.022 | 143 |
| Right Somatosensory Cortex | 38, -12, 20 | 1 | 4.50 | <.001 | 385 |
| Left Somatosensory Cortex | -42, -20, 20 | 1 | 4.40 | <.001 | 280 |
| Right Posterior Cingulate Cortex (PCC) | 8, -44, 30 | 23 | 3.97 | <.001 | 335 |
| *Schizophrenia > Healthy Controls* |  |  |  |  |  |
| Prefrontal Cortex/Response Inhibition | 28, 48, 26 | 9 | 4.47 | 0.013 | 159 |
| Left Frontal Eye Fields (FEF) | -28, -6, 46 | 6 | 4.65 | 0.020 | 146 |
| Right Cerebellum (Ventral Attention)<sup>b</sup> | 40, -54, -32 | N/A | 5.80 | <.001 | 466 |
| Left Cerebellum (Ventral Attention)<sup>b</sup> | -32, -56, -28 | N/A | 5.87 | <.001 | 564 |
| Left Cerebellum (Ventral Attention)<sup>b</sup> | -2, -66, -20 | N/A | 4.25 | 0.003 | 207 |
| Precuneus | 8, -68, 52 | 7 | 4.20 | 0.001 | 276 |
| Ventral Posterior Cingulate Cortex (PCC) | 0, -38, -2 | N/A | 4.67 | 0.042 | 122 |
| Right Parietal | 42 -44 56 | 7 | 4.79 | 0.013 | 161 |
| Left Superior Parietal Cortex | -16 -64 60 | 7 | 4.23 | <.001 | 282 |

<sup>a</sup>Values represent family-wise error corrected rate; <sup>b</sup>Cerebellum coordinates were compared to cerebellar parcellations provided in Buckner et al., (2011), *Journal of neurophysiology*. Network names in parentheses indicate the network that that region of the cerebellum is primarily functionally connected with, based on the Buckner parcellation.

**Fig. 1:**
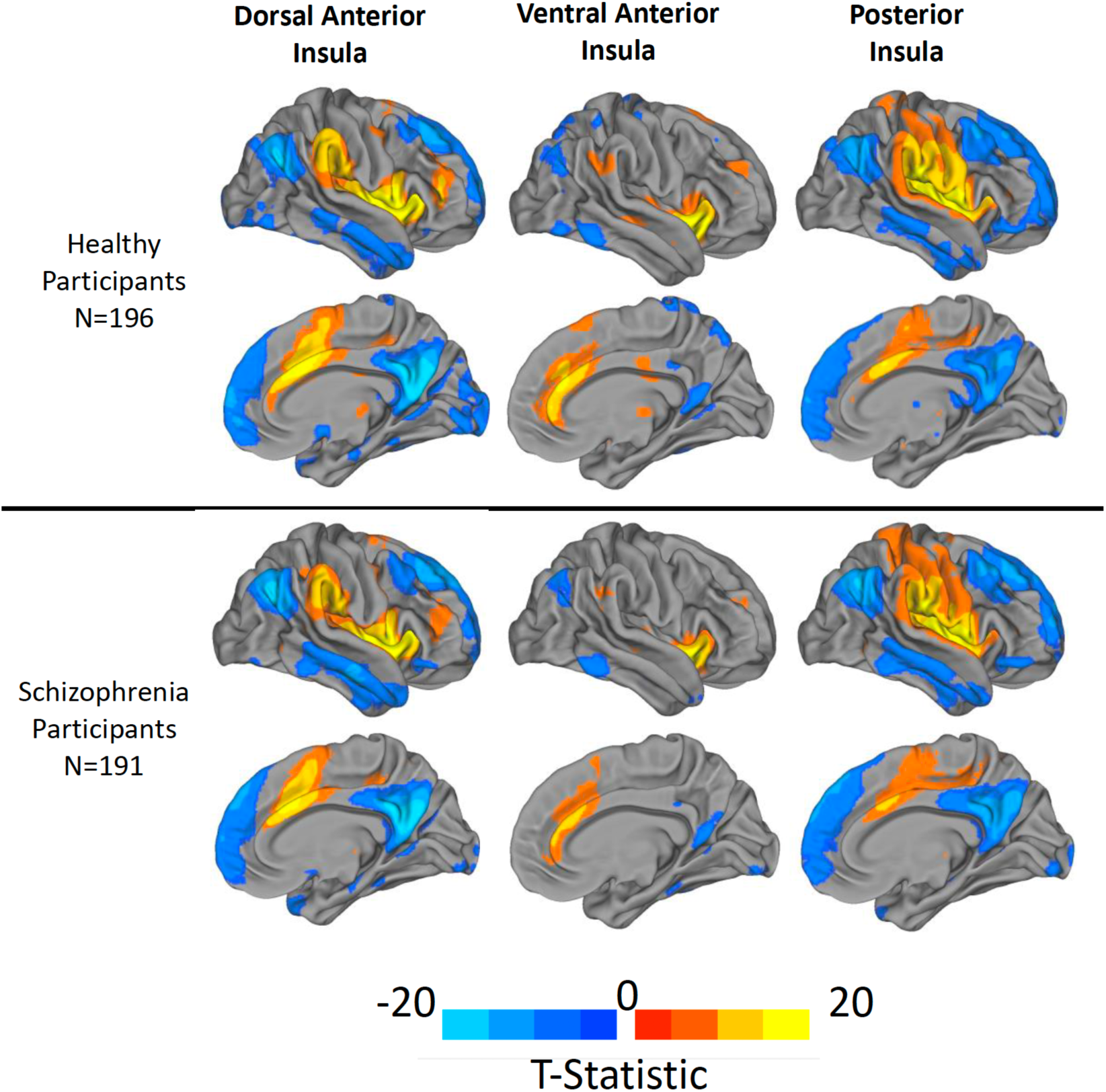
Within-group functional connectivity of insula sub-regions projected onto inflated right-hemisphere surfaces to illustrate functional profiles of each sub-region. Results thresholded at voxel-wise pFWE<.05.

**Fig. 2:**
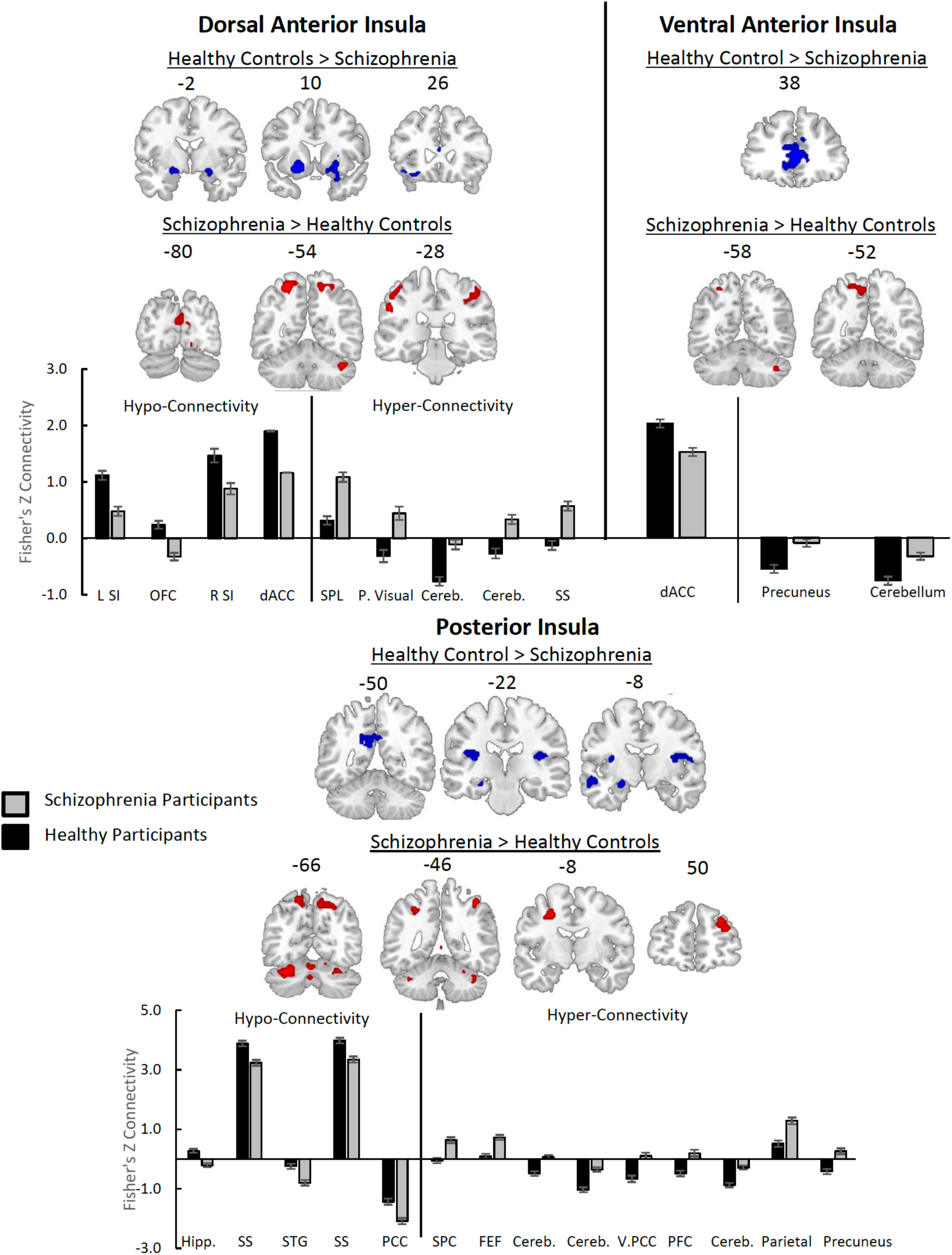
Clusters demonstrating significant hypoconnectivity (healthy > schizophrenia) and hyperconnectivity (schizophrenia > healthy) for each insula sub-region. Group difference maps were thresholded at cluster-level p_FWE_ <.05 for voxel-wise cluster-defining threshold p=.001 (uncorrected), controlling for study protocol and gender. Bar plots illustrate average functional connectivity within each significant cluster. *L SI = Left Substantia Innominata; OFC = Orbitofrontal Cortex; R SI = Right Substantia Innominata; SPL = Superior Parietal Lobe; P. Visual = Primary Visual Cortex; SS = Somatosensory Cortex; dACC = Dorsal Anterior Cingulate Cortex; Hipp. = Hippocampus; STG = Superior Temporal Gyrus; PI = Posterior Insula; PCC = Posterior Cingulate Cortex; SPC = Superior Parietal Cortex; FEF = Frontal Eye Field; Cereb. = Cerebellum; V. PCC = Ventral Posterior Cingulate Cortex; PFC = Prefrontal Cortex.*

#### 3.2.1 Dorsal Anterior Insula (dAI)

Schizophrenia was characterized by a combination of dAI hypo- and hyper-connectivity. Specifically, dAI positive connectivity with bilateral sublenticular substantia innominata (part of the extended amygdala), dACC, and the orbital frontal cortex (OFC) was reduced in schizophrenia (i.e. hypo-connectivity). In contrast, dAI connectivity with sensorimotor and attention regions was increased in schizophrenia (i.e. hyper-connectivity). Relatively stronger positive connectivity between dAI and left parietal cortex, primary visual cortex, right somatosensory cortex, and the cerebellum was observed in schizophrenia as compared to healthy participants. Additionally, dAI was negatively connected with a region in the cerebellum associated with the ventral attention network in healthy participants (Buckner et al., 2011) but exhibited minimal connectivity (close to zero) in the schizophrenia group.

#### 3.2.2. Ventral Anterior Insula (vAI)

Reduced positive connectivity of the vAI and dACC were seen in schizophrenia. Furthermore, vAI was more weakly negatively connected with regions comprising the DMN, namely the precuneus and cerebellum.

#### 3.2.3 Posterior Insula (PI)

Individuals with schizophrenia demonstrated widespread dysconnectivity within the PI. Reduced positive connectivity in schizophrenia was seen between PI and the superior temporal gyrus, hippocampus, and somatosensory cortex, relative to the connectivity observed in healthy participants. The PI was negatively connected with the PCC in both groups, but the strength of that negative connectivity was weaker in the schizophrenia participants.

In schizophrenia, the PI was more strongly positively connected to the superior parietal lobe, frontal eye fields, and right parietal cortex – regions that exhibited minimal connectivity in healthy participants (i.e. connectivity close to zero). There were also several regions with strong negative connectivity in the healthy participants that showed weaker negative connectivity in schizophrenia. These regions included the prefrontal cortex, precuneus, regions of the cerebellum associated with the ventral attention network, and a ventral portion of the PCC.

#### 3.2.4 Similarity Analysis

As a complementary analysis to the above findings, similarity between insula sub-region connectivity maps was computed using eta-squared and compared between groups (Table S3). PI-dAI connectivity profiles were significantly more similar in schizophrenia than healthy participants (eta-squared 0.023 higher, 95% CI(0.008, 0.04)) whereas dAI-vAI connectivity profiles were significantly less similar (eta-squared 0.043 lower, 95% CI(−0.079, −0.005)). The similarity between vAI-PI maps did not differ between groups (eta-squared 0.021 lower in schizophrenia, 95% CI(−0.061, 0.031)). These findings further demonstrate a greater similarity in functional profiles between the PI and dAI and less similar functional profiles between the dAI and vAI in schizophrenia than is typical.

### 3.3 Associations Between Insula Connectivity and Clinical Variables

#### 3.3.1 Region of Interest Correlation Analysis

Consistent with our hypothesis, dAI hypo-connectivity correlated with cognitive function (r=.18, p=.014), surviving multiple comparisons correction (Fig. 3a). Neither positive nor negative symptoms correlated with average dAI hypo-connectivity (positive: r=-.002, p=.978; negative: r=.01, p=.949), suggesting a specific relationship between dAI connectivity and cognition. Our hypotheses that vAI hypo-connectivity would correlate with negative symptoms (r=-.01, p=.944) and PI hyper-connectivity would correlate with positive symptoms (r=-.05, p=.538) were not supported.

**Fig. 3:**
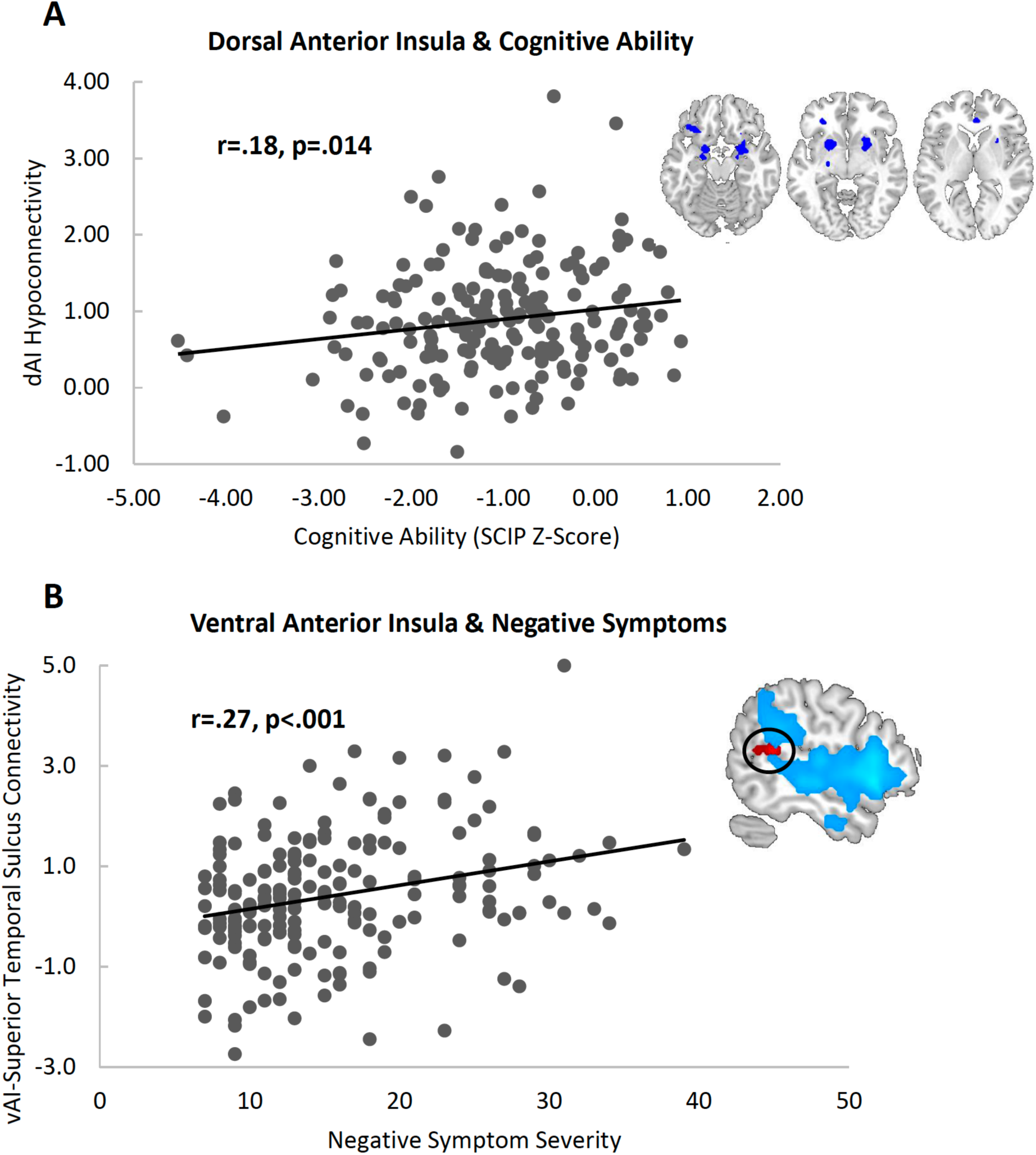
A) Average connectivity extracted from the regions that demonstrated reduced connectivity with the dAI in schizophrenia (shown in blue) was associated with better overall cognitive ability (total SCIP z-score). B) Whole-brain voxel-wise analysis revealed a positive association between vAI connectivity with right posterior superior temporal gyrus (shown in red) and negative symptoms. Connectivity of vAI with the whole brain in the healthy group is overlaid in blue, demonstrating no significant connectivity between vAI and STS in healthy participants, suggesting that vAI-STS are abnormally connected in schizophrenia, contributing to negative symptom severity.

#### 3.3.2 Whole Brain Correlation Analysis

In an effort to further explore brain-behavior relationships between insula sub-regions and clinical characteristics of schizophrenia, whole brain voxel-wise correlations were performed for sub-region correlations and specific *a priori* symptoms. At cluster-level p_FWE_<.05 threshold, connectivity between vAI-posterior superior temporal sulcus (STS) was positively correlated with negative symptom severity (r=.27, p<.001) (Fig. 3b). Relationships between dAI and cognition and between PI and positive symptoms were not significant at this threshold.

## 4. Discussion

In contrast to well-documented structural abnormalities of the insula, comparatively little is known about functional connectivity disturbances of the insula in schizophrenia. This is especially true for insula sub-regions, including dAI, vAI, and PI divisions, which demonstrate distinct connectivity profiles in keeping with their divergent functional roles. To address this knowledge gap, we analyzed functional connectivity of insula sub-regions in a relatively large cohort of people with schizophrenia and healthy individuals. We found widespread insula dysconnectivity in schizophrenia characterized by both hypo- and hyper-connectivity of all three insula sub-regions. Further comparison revealed greater similarity between dAI and PI functional profiles in schizophrenia than healthy participants, as well as less similarity between dAI and vAI. DAI hypoconnectivity correlated with cognitive impairment and vAI-STS hyper-connectivity correlated with negative symptoms. Our findings add to prior literature (Tian et al., 2019) suggesting a less differentiated insula connectome in schizophrenia that is associated with clinical phenotypes.

Only one prior study of insula connectivity in schizophrenia has examined insula sub-regions, yielding some convergent findings (Chen et al., 2016). In a smaller, older sample of patients, Chen and colleagues (Chen et al., 2016) also observed dAI dysconnectivity with OFC, the basal ganglia, and dACC. However, they observed no differences in vAI connectivity and their main finding of increased PI-thalamus connectivity was not replicated here. Recent analysis of anterior/posterior insula connectivity also found reduced anterior insula connectivity with OFC, ACC, and basal ganglia, as well as reduced PI connectivity with superior temporal gyrus and PCC (Tian et al., 2019). Our results build on these previous findings and, in a large, relatively young sample, characterize insula functional connectivity in schizophrenia in the following ways: 1) all three sub-regions are hypo-connected with areas typically connected in healthy participants. This follows the structural literature identifying widespread structural abnormalities of the insula in schizophrenia (e.g. reduced volume and thickness), suggesting that the entire insula is impacted (Shepherd et al., 2012); 2) dAI and vAI are *less* strongly connected with DMN, salience, and central executive regions while PI is *more* strongly negatively connected with the PCC, a hub of the DMN. Connectivity abnormalities between DMN, salience and central executive networks is theorized to be a core contributor to the pathophysiology of schizophrenia (Menon, 2011); 3) dAI is hyperconnected with somatosensory/parietal regions and PI is hyperconnected with cognitive regions, indicating inappropriate connectivity of dAI with regions typically connected to PI, and vice versa. This atypical connectivity pattern was further quantified in a similarity analysis, which showed that the connectivity profiles of these two sub-regions were significantly more similar in schizophrenia than healthy participants, suggesting less clearly differentiated functional profiles. Recently, Tian and colleagues (Tian et al., 2019) reported reduced differentiation of anterior/posterior insula connectivity in schizophrenia based on reduced modularity (Q) of insula clusters and a more diverse connectivity profile along the rostrocaudal axis. Our findings further suggest that dAI and PI sub-networks, which typically support different aspects of information processing (i.e. cognitive and somatosensory), are less specifically connected in schizophrenia, leading to a more disorganized insula connectome.

Motivated by a growing awareness of the functions of insula sub-regions, we tested several a-priori hypotheses on the associations between insula functional dysconnectivity in schizophrenia. As briefly reviewed earlier, our hypotheses were guided by evidence that dAI, vAI, and PI are preferentially involved in cognition, emotion processing, and somatosensory functions, respectively. In line with our first hypothesis, the overall reduction observed in dAI connectivity was associated with greater cognitive impairment, which is consistent with prior findings implicating dAI in cognitive ability (Moran et al., 2013; Sheffield et al., 2015). Our second hypothesis, that vAI hypo-connectivity would be associated with more severe negative symptoms, was not supported. However, whole-brain voxel-wise analysis within the schizophrenia group identified *stronger* connectivity between vAI-STS related to worse negative symptoms. The posterior STS is considered a core area involved in social cognition (Deen et al., 2015) that exhibits hyperactivity during processing of neutral faces in schizophrenia (Mier et al., 2017) and is implicated in the social impairments observed in autism (Zilbovicius et al., 2006). The vAI also plays a major role in social cognition through emotional understanding (Gallese et al., 2004) and therefore stronger vAI-STS connectivity could contribute to inappropriate processing of emotion during the viewing of neutral faces, leading to social cognitive deficits. While the insula and STS are both implicated in models of social cognitive deficits in schizophrenia (Billeke and Aboitiz, 2013) these are some of the first findings to demonstrate that their connectivity is associated with negative symptom severity in schizophrenia. Our third hypothesis of PI hyper-connectivity correlating with severity of positive symptoms was also not supported, in either the ROI or whole-brain analysis. It is possible that resting-state connectivity is not the best way to elucidate associations with psychotic experiences. More direct analysis of PI functioning in schizophrenia, such as was seen in symptom-capture studies (Mallikarjun et al., 2018), may be more sensitive to relationships between PI integrity and psychosis. Finally, all brain-behavior relationships were hypothesized based on canonical (i.e. “healthy”) sub-region functional profiles. We found reduced specificity of these functional profiles in schizophrenia. Therefore, while speculative, less differentiated functional profiles may have made it more difficult to detect the expected brain-behavior associations for all three insula sub-regions.

The general finding of reduced functional differentiation of the insula suggests a more disorganized insula connectome in schizophrenia, thereby implicating developmental processes in these abnormal connectivity profiles. The insula is the earliest cortical region to develop and differentiate, starting at 6 weeks of fetal life (Chi et al., 1977) and the anterior-posterior gradient is one of the first identifiable factors of embryonic development (Fjell et al., 2018). Longitudinal imaging of infants has revealed anterior/posterior functional sub-divisions within the first two years of life. At 33 days, the anterior and posterior insula show diffuse connectivity with frontal and somatosensory regions, respectively, indicating very early formation of distinct functional profiles (Alcauter et al., 2015). By two years, these sub-regions exhibit network-like connectivity with distributed brain regions. Insula connectivity continues to be refined throughout childhood and adolescence (Uddin et al., 2011) and reduced centrality of the anterior insula is associated with childhood maltreatment (Teicher et al., 2014), indicating that its connectivity may be particularly vulnerable to developmental insult. Interestingly, in our data, the anterior sub-regions were inappropriately connected with more posterior brain areas while the PI was inappropriately connected with more anterior brain areas, reflected in greater similarity of functional profiles. The vAI and dAI were also *less* similar in schizophrenia, suggesting that connectivity from anterior insula was being “pulled away” to posterior insula. As described above, the insula undergoes multifaceted changes in connectivity throughout development, providing many opportunities for insula specificity to go awry. Schizophrenia has been associated with both pre- and perinatal risk factors such as viral infections and stress of the mother during pregnancy (King et al., 2010), childhood maltreatment (DeRosse et al., 2014) and environmental stress (van Os et al., 2010). Abnormal connectivity of all three insula sub-regions converges with the notion that altered insula structure and function may be shaped throughout development in ways that confer risk for schizophrenia and, although speculative, suggests that the specific connections associated with insula sub-regions may develop in a more disorganized fashion, contributing to the pathophysiology of schizophrenia.

Regarding limitations, nearly all schizophrenia participants were taking anti-psychotic medications. The vast majority of clusters demonstrating group differences in connectivity were unrelated to medication dosage suggesting that the widespread alterations in connectivity observed in our data are unlikely to be driven by medication use. Additionally, in an effort to stick closely with our *a priori* brain-behavior relationships and limit exploratory correlations, associations between insula sub-region connectivity and clinical characteristics that could have reasonably been hypothesized (e.g. dAI and positive symptoms (Palaniyappan and Liddle, 2012)) were not tested. Finally, although positive and negative symptoms were assessed very close to the time of the scan, the PANSS is still a fairly distal measure of symptoms, making it more difficult to detect brain-behavior relationships.

In conclusion, there is significant and widespread dysconnectivity of insula sub-regions in schizophrenia. This dysconnectivity is characterized by hypo-connectivity with regions typically connected in the healthy brain, altered connectivity with DMN regions, and inappropriate connectivity of dAI with parietal/somatosensory regions and PI with higher-order cognitive regions, revealing a disorganized connectome. Hypoconnectivity of dAI was associated with worse cognition, while vAI-STS connectivity related to negative symptoms. It is clear from these results that the well-characterized structural abnormalities of the insula in schizophrenia have consequences for functional connectivity compelling future research into structure and function of insula sub-regions throughout development and illness course.

## Supplemental Materials

### Methods

#### Participant Exclusion Criteria

Exclusion criteria were nearly identical across the three neuroimaging studies and included age less than 16 or older than 65 (age 55 in the case of 1R01MH102266); history of significant head trauma, medical illness or central nervous system disorder (e.g. epilepsy); substance abuse within the last one month for schizophrenia participants (three months in the case of 1R01MH102266) or lifetime history of substance abuse/dependence in healthy controls; estimated premorbid IQ less than 70 based on the Wechsler Test of Adult Reading (WTAR)(Wechsler, 2001); MRI contra-indicators; healthy subjects could not have a first degree relative with a psychotic illness.

Of the 444 individuals who participated in one of the three MRI studies, 57 (14 healthy, 43 schizophrenia) were excluded for one of the following reasons: not meeting fMRI quality assurance (QA) threshold as described below (six healthy, 30 schizophrenia), resting-state scan was not acquired during scanning session (three healthy, four schizophrenia), presence of obvious artifacts on T1-weighted scan (three healthy, two schizophrenia), or segmentation failed (two healthy, seven schizophrenia).

#### Scanning Parameters

Imaging data were collected on two identical 3T Philips Intera Achieva MRI Scanners located at the Vanderbilt University Institute of Imaging Science. Parameters differed slightly across studies. For R01MH070560, a 7-minute echo-planar imaging resting-state scan was acquired with 203 volumes, 90 degree flip angle, FOV = 80×80, 3.0×3.0×3.0mm resolution, slice thickness=4.0mm, TR/TE=2000/35ms. For CT00762866, 7-minute resting-state scans were collected with 203 volumes, 90 degree flip angle, FOV = 80×80, 3.0×3.0×3.2mm resolution, 3.2mm slice thickness, TR/TE=2000/35ms. Both studies also acquired T1-weighted fast field echo (FFE) structural scans (170 sagittal slices, FOV = 256×256, 1.0mm isovoxel resolution, TR/TE=8.0/3.7ms). Data collected as part of R01MH102266 included a 10-minute resting-state scan with 300 volumes, FOV = 80×80, 3.0×3.0×3.0mm resolution, slice thickness=3.3mm, TR/TE=2000/43ms and a T1-weighted structural scan (170 sagittal slices, FOV = 256×256, 1.0mm isovoxel resolution, TR/TE=8.9/3.7ms). In all studies, subjects were instructed to remain awake and rest quietly with their eyes closed.

#### Preprocessing

The processing pipeline was containerized using Singularity, built at SingularityHub ((Sochat et al., 2017) https://singularity-hub.org), and executed using the Vanderbilt University Institute of Imaging Science XNAT infrastructure (Harrigan et al., 2016). T1-weighted anatomical images were segmented into grey matter, white matter, and cerebrospinal fluid (CSF) partitions using a multi-atlas algorithm as previously described (Asman and Landman, 2013; Sheffield et al., 2019). The grey matter image was then nonlinearly transformed (Ashburner & Friston, K. J., 1999; Ashburner Neelin, P., Collins, D. L., Evans, A., & Friston, K., 1997) to align with the MNI space gray matter template provided with SPM12 ((Wellcome Centre for Human Neuroimaging, https://www.fil.ion.ucl.ac.uk/spm/software/spm12).

#### QA Procedure

Neuroimaging data underwent quality assurance (QA) as described previously by our group (see (Sheffield et al., 2019)). Briefly, T1-weighted and EPI functional data were first visually inspected for obvious neuroimaging artifacts and segmentation/normalization failures. Median voxel displacement log10 and median temporal signal-to-noise (SNR) was then calculated for each subject’s fMRI time-series data. Scans with median temporal signal-to-noise ratios below the 5th percentile or median intraframe voxel displacements above the 95th percentile of the entire sample were considered to have excessive head motion and were excluded from further analysis.

## Group Differences in Sub-region Connectivity for Left and Right Insula

**Supplemental Table 1:**
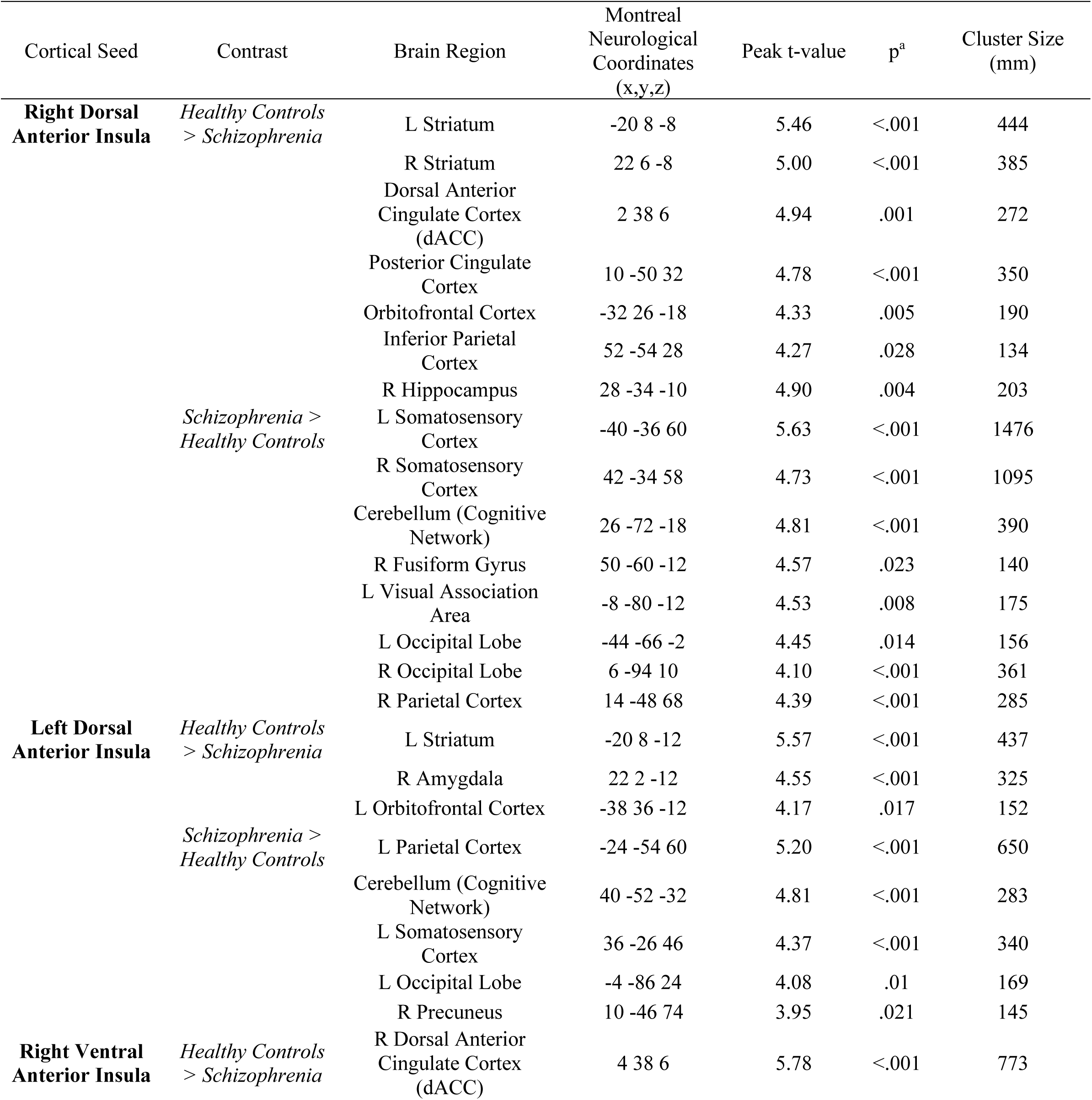
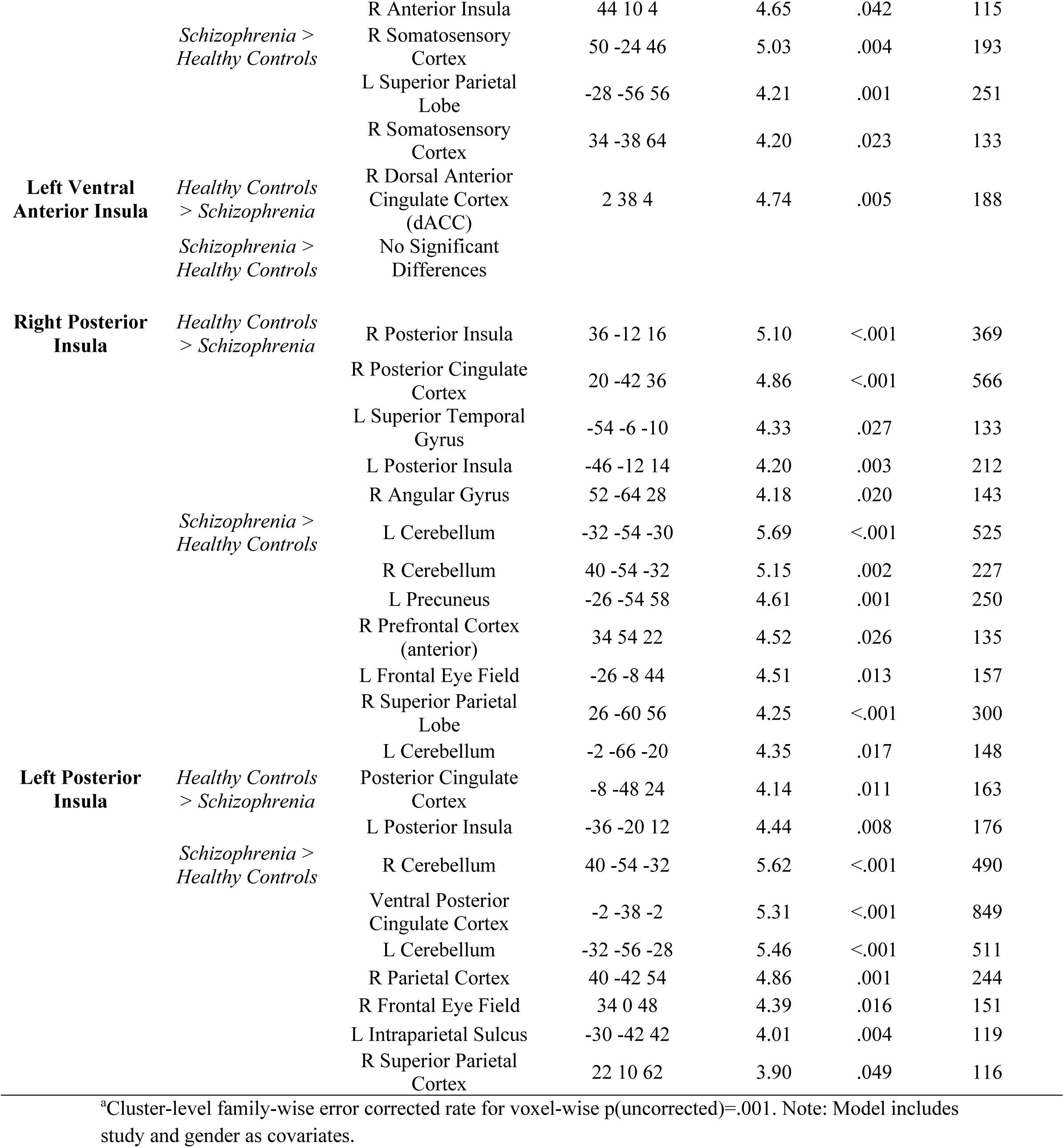
Insula functional connectivity abnormalities in schizophrenia

**Supplemental Table 2:** Associations between clusters showing significant group differences in connectivity and chlorpromazine-equivalent dose, in schizophrenia participants

|  | **MNI Coordinates**<br>**X, Y, Z** | **Correlation with CPZ-**<br>**Equivalent Dose (N=165)** |
| --- | --- | --- |
| **Dorsal Anterior Insula** |  |  |
| L. Substantia Innominata | -20, 8, -10 | $r=-.08, p=.296$ |
| OFC | -30, 36, -8 | $r=.04, p=.616$ |
| R. Substantia Innominata | 22, 2, -12 | $r=-.14, p=.078$ |
| dACC | 2, 38, 6 | $r=-.01, p=.892$ |
| Superior Parietal Lobe | -24, -54, 60 | $r=-.06, p=.441$ |
| Primary Visual Cortex | 22, -68, 10 | $r=-.10, p=.194$ |
| Occipital Lobe | 26, -72, -20 | $r=-.05, p=.510$ |
| Cerebellum | 40, -52, -32 | $r=-.01, p=.928$ |
| Somatosensory Cortex | 42, -34, 58 | $r=-.05, p=.569$ |
| **Ventral Anterior Insula** |  |  |
| dACC | 2, 38, 6 | $r=.02, p=.832$ |
| Precuneus | -8, -54, 52 | $r=-.01, p=.890$ |
| Cerebellum | 40, -60, -38 | $r=-.02, p=.834$ |
| **Posterior Insula** |  |  |
| Hippocampus | -24, -10, -12 | $r=.01, p=.935$ |
| Somatosensory Cortex | -42, -20, 20 | $r=-.19, p=.014$ |
| Superior Temporal Gyrus | -56, -6, -10 | $r=-.03, p=.663$ |
| Somatosensory Cortex | 38, -12, 20 | $r=-.27, p<.001$ |
| PCC | [8 -44 30] | $r=.06, p=.464$ |
| Superior Parietal Cortex | -16, -64, 60 | $r=-.04, p=.650$ |
| Frontal Eye Fields | -28, -6, 46 | $r=-.08, p=.300$ |
| Cerebellum | -2, -22, -20 | $r=.02, p=.810$ |
| Cerebellum | -32, -56, -28 | $r=.08, p=.336$ |
| Ventral PCC | 0, -38, -2 | $r=-.06, p=.452$ |
| Prefrontal Cortex | 28, 48, 26 | $r=-.05, p=.494$ |
| Cerebellum | 40, -54, -32 | $r=.06, p=.422$ |
| Parietal Lobe | 42, -44, 56 | $r=-.01, p=.888$ |
| Precuneus | 8, -68, 52 | $r=-.03, p=.692$ |

### Similarity Analysis

Significant group differences in eta-squared reflected whether insula sub-networks were more or less similar to one another (i.e. differentiated) in schizophrenia compared to healthy participants. Empirical eta-squared values were compared between groups using a bootstrap distribution (300 bootstrap samples) to provide 95% percentile confidence intervals (CI) for each group difference.

**Supplemental Table 3.**
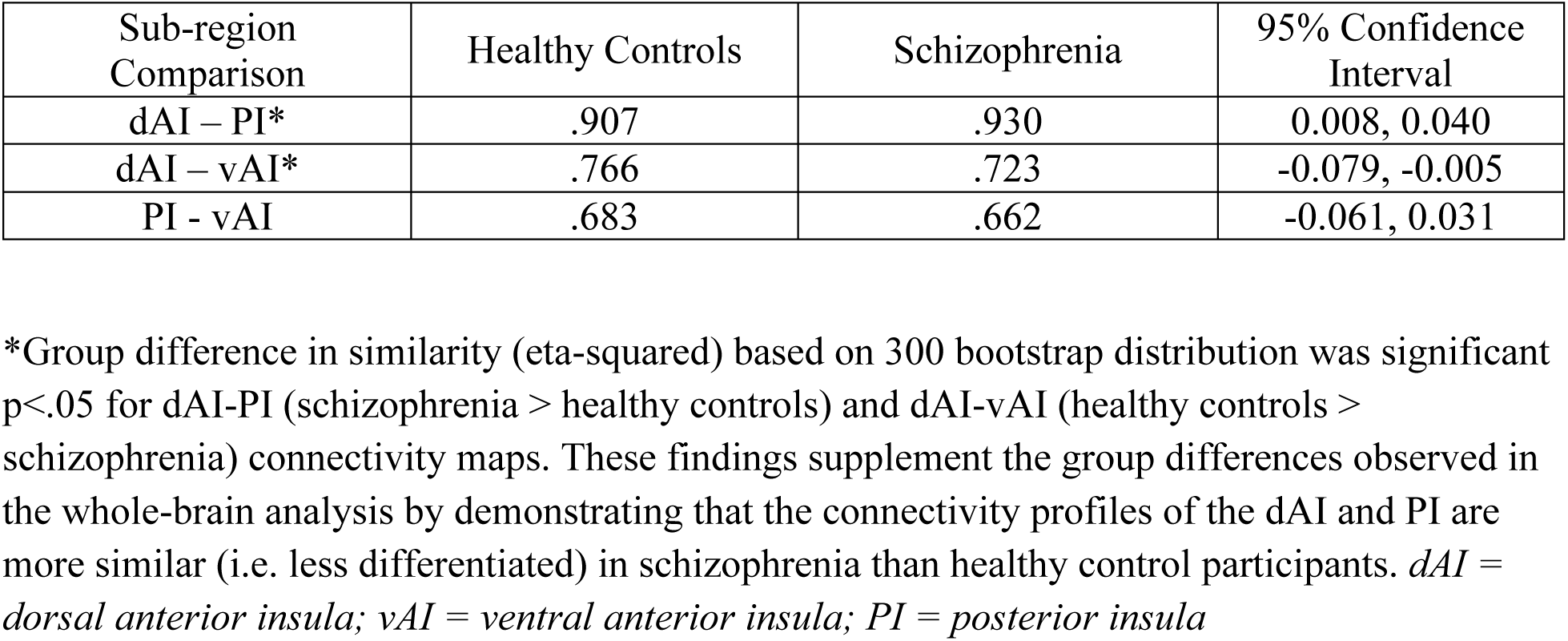

